# Snail shell colour evolution in urban heat islands detected via citizen science

**DOI:** 10.1101/424564

**Authors:** Niels A. G. Kerstes, Thijmen Breeschoten, Vincent Kalkman, Menno Schilthuizen

## Abstract

The extreme environmental conditions that prevail in cities are known to cause selection pressures leading to adaptive changes in wild, city-dwelling, organisms (“urban evolution”). The urban heat island, elevated temperatures in the city centre due to a combination of generation, reflection, and trapping of heat, is one of the best recognised and most widespread urban environmental factors. Here, we used a citizen-science approach to study the effects of urban heat on genetically-determined shell colour in the land snail *Cepaea nemoralis* in the Netherlands. We used smartphone applications to obtain colour data on almost 8,000 snails throughout the country. Our analysis shows that snails in urban centres are more likely to be yellow than pink, an effect predicted on the basis of thermal selection. Urban yellow snails are also more likely to carry dark bands at the underside of the shell; these bands might affect thermoregulation in yet underexplored ways.

## Introduction

It is increasingly understood that the environmental changes that humans are causing in a growing part of the earth’s surface, especially in heavily urbanised areas, are becoming a major evolutionary force for the organisms inhabiting those places^1,9^. Numerous examples exist of such “urban evolution”, e.g., species of city-dwelling animals and plants adapting to urban microclimate, urban habitat fragmentation, and urban pollution^11,23^.

Urban evolution is a process that, by definition, takes place in areas densely populated by people. Moreover, since certain aspects of the urban environment (such as the “urban heat island”^4^) are shared by cities worldwide, different cities could be viewed as replicates for studying specific evolutionary responses in widespread species^10^. For these reasons, urban evolution is eminently suited to be studied through the dispersed monitoring systems that citizen science provides. However, although the 2009 “Evolution Megalab” (www.evolutionmegalab.org; Ref. 27) has shown that citizen science methods enable the tracking of non-urban contemporary evolutionary change, urban evolution has not yet been targeted by any major citizen science projects.

To make a citizen science approach to urban evolution possible, several conditions need to be met^26,29^. First of all, the organism of study must be widespread, harmless, common, large enough to be observed and studied without the need of specialised equipment, and carry externally visible genetic variation that responds to a universal urban selection pressure. Second, a simple digital data collection platform needs to be present to enable a large number of untrained citizen scientists to upload data. Finally, some pre-existing knowledge on the evolutionary ecology of the species in question is required for hypotheses to be formulated and tested.

In this paper, we describe such a citizen science approach to urban evolution in the Netherlands. We developed a simple smartphone app (‘SnailSnap’) for taking and uploading images of the shell colour polymorphism in ***Cepaea nemoralis***, an intensively-studied land snail that has been a model species in evolutionary ecology and genetics for over a hundred years^12,15^. The app was linked to a popular Dutch citizen science platform, which allowed us to test the hypothesis that thermal selection of the urban heat island results in an evolutionary response in genetically determined snail shell colour.

## Results

The ‘SnailSnap’ app was downloaded 1,180 times, and 9,483 images of ***C. nemoralis*** were uploaded. After removal of pictures that could not be unequivocally categorized (see Methods), we retained 7,868 images of unique, individual snails and investigated the relationship between genetically-based shell colour and four habitat types: [1] agricultural land, [2] nature, including forests, [3] urban green areas, i.e., urban parks and forests, sport and recreational areas, and [4] urban “grey” areas, i.e., residential, commercial, and industrial areas.

### Colour

We found that the proportion of yellow snails is higher in urban than in non-urban areas (Fig. 1, χ^2^ = 12.35, p = 0.0004), while the proportion of pink snails is lower (χ^2^ = 14.21, p = 0.0002). The proportions of brown snails did not differ significantly (χ^2^ = 1.96, p = 0.58). There was no significant difference in the proportion of yellow snails between urban grey and urban green areas (χ^2^ = 0.08, p = 0.77). The proportion of yellow three-banded snails in the urban grey habitat was higher than in agricultural (χ^2^ = 8.78, p = 0.003), natural (χ^2^ = 12.91, p = 0.0003), or urban green areas (χ^2^ = 4.78, p = 0.029).

**Figure 1.**
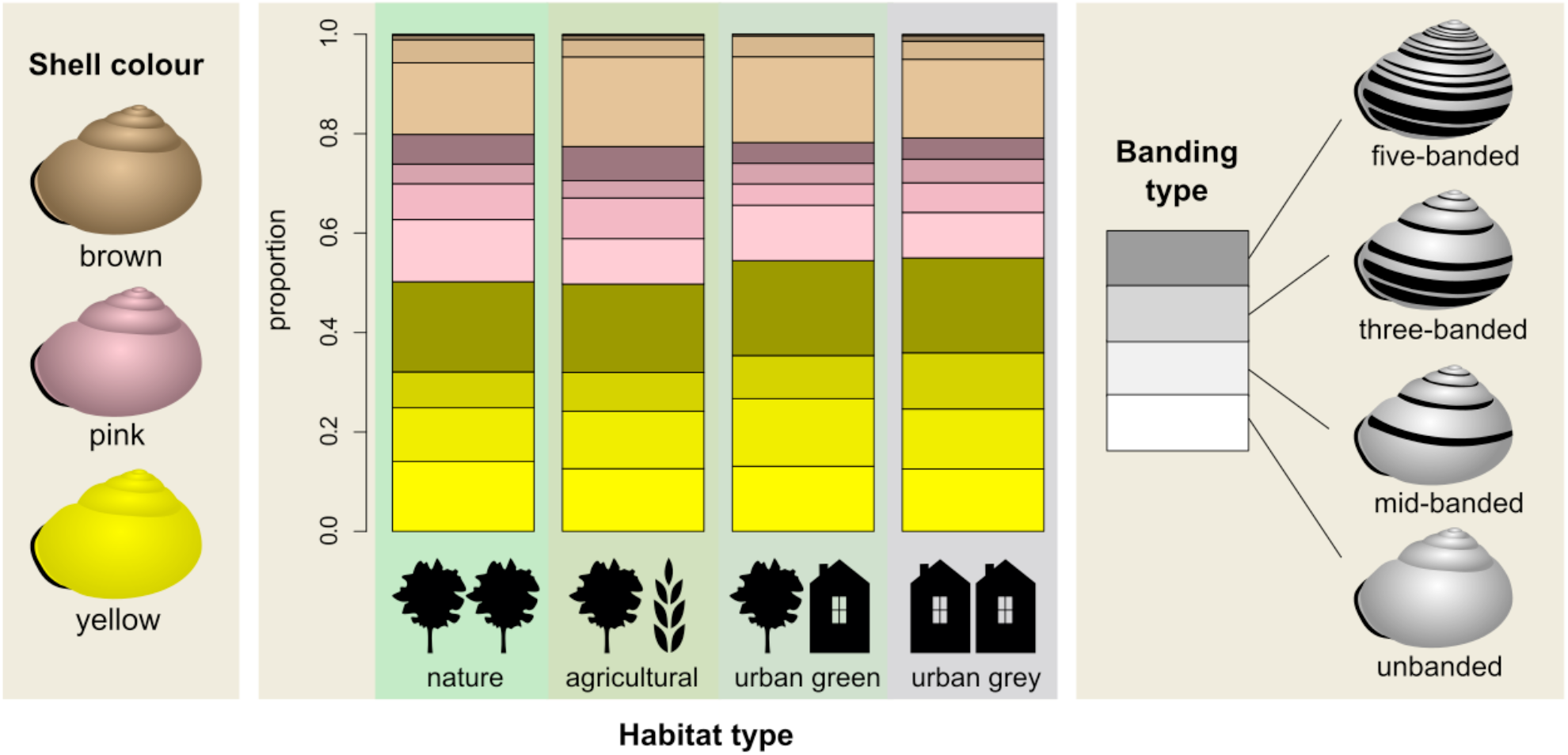
Proportions of shell colour (yellow, pink or brown) and the four main banding types per habitat type.

The results of the multinomial logistic regression with shell colour as outcome variable (table 1) provide further explanation for these patterns. Yellow was chosen as the reference colour. When including snails from all habitat types, the probability of a pink snail decreases with increasing temperature relative to the probability of a yellow snail (Odds-ratios < 1). The effect of temperature on the probability of a brown snail depends on the number of dry days; at sites with a low number of dry days, the probability of a brown snail decreases with increasing temperature. At sites with a high number of dry days, the proportion of brown snails is very low overall and the effect of temperature on the proportion of brown snails is negligible.

**Table 1.**
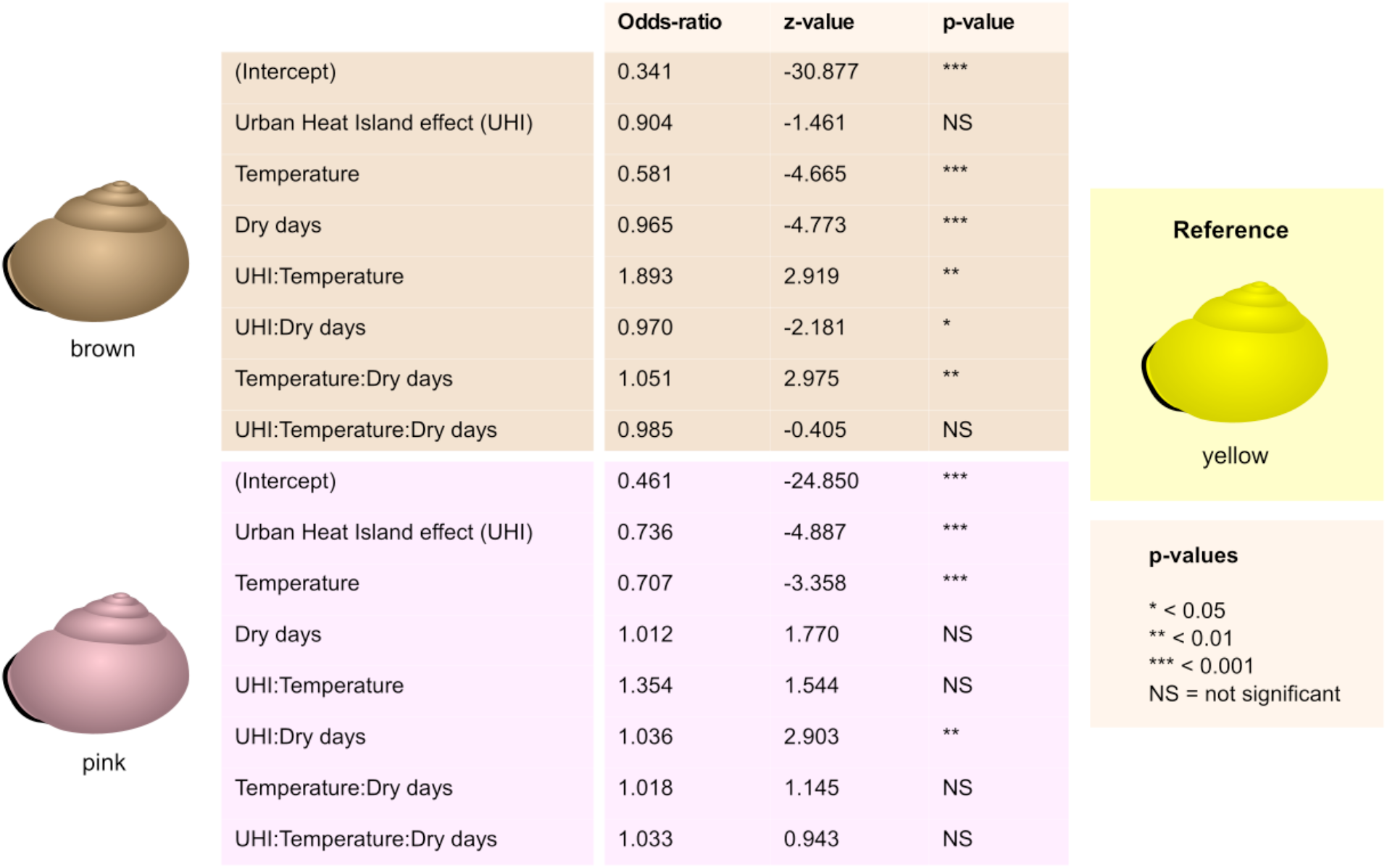
Results of a multinomial regression model with colour as outcome variable, including snails from all land use types (n = 7,868). Yellow snails are the reference.

|  |  | Odds-ratio | z-value | p-value |  |
| --- | --- | --- | --- | --- | --- |
| brown | (Intercept) | 0.341 | -30.877 | *** | Reference<br>yellow |
|  | Urban Heat Island effect (UHI) | 0.904 | -1.461 | NS |  |
|  | Temperature | 0.581 | -4.665 | *** |  |
|  | Dry days | 0.965 | -4.773 | *** |  |
|  | UHI:Temperature | 1.893 | 2.919 | ** |  |
|  | UHI:Dry days | 0.970 | -2.181 | * |  |
|  | Temperature:Dry days | 1.051 | 2.975 | ** |  |
|  | UHI:Temperature:Dry days | 0.985 | -0.405 | NS |  |
| pink | (Intercept) | 0.461 | -24.850 | *** | p-values<br>* < 0.05<br>** < 0.01<br>*** < 0.001<br>NS = not significant |
|  | Urban Heat Island effect (UHI) | 0.736 | -4.887 | *** |  |
|  | Temperature | 0.707 | -3.358 | *** |  |
|  | Dry days | 1.012 | 1.770 | NS |  |
|  | UHI:Temperature | 1.354 | 1.544 | NS |  |
|  | UHI:Dry days | 1.036 | 2.903 | ** |  |
|  | Temperature:Dry days | 1.018 | 1.145 | NS |  |
|  | UHI:Temperature:Dry days | 1.033 | 0.943 | NS |  |

UHI (Urban Heat Island effect) is a significant predictor for the probability of a pink versus a yellow snail. There is also a significant interaction between UHI and the number of dry days for the probability of a pink versus a yellow snail. At sites with relatively low numbers of dry days, the proportion of pink snails decreases with increasing UHI. At sites with a relatively high number of dry days, UHI does not result in a decrease in the probability of pink snails. For the probability of a brown snail, there is a significant and fairly strong interaction between UHI and temperature (figure 2). Furthermore, the probability of a brown snail decreases with an increasing number of dry days (figure S1). When only snails from urban grey areas are included, we still find a relationship between UHI and shell colour (table S1).

**Figure 2.**
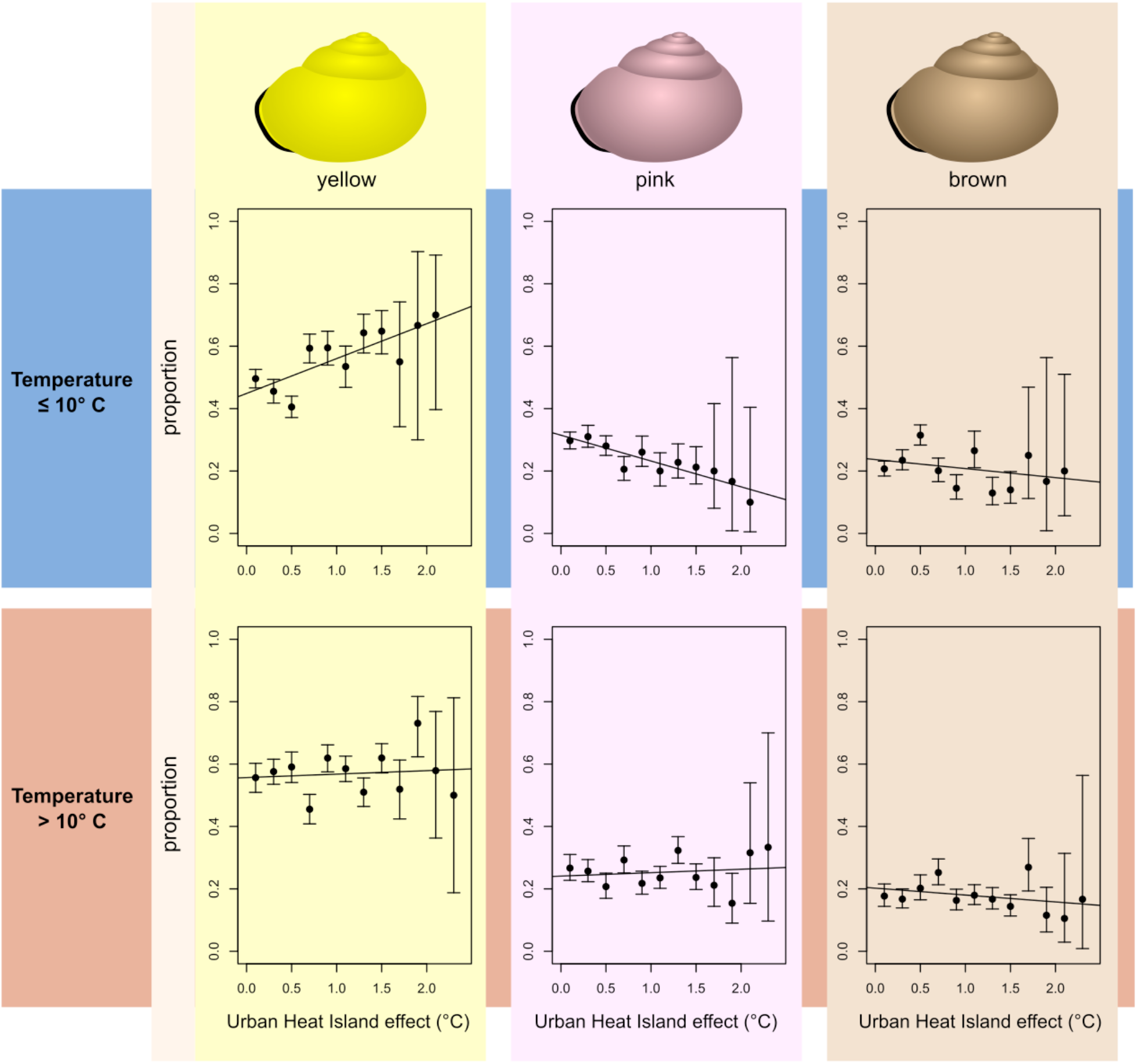
The proportions of yellow, pink and brown snails as a function of the Urban Heat Island effect (UHI). Error bars indicate 95% confidence intervals (Wilson score interval). Based on the median average temperature (10°C), the data set was split into a “cold” half (A, n = 3,936) and a “hot” half (B, n = 3,932). Starting at UHI = 0°C, UHI was divided into categories of 0.2°C and proportions per category were plotted against the central value of each category. Linear regression lines are included for illustrative purposes.

### Banding

Since brown snails are rarely banded, multinomial regression models with banding type as outcome variable were only performed for yellow and pink snails (table 2). “Unbanded” was chosen as the baseline banding type. The probabilities of unbanded and five-banded yellow snails decrease with increasing temperature, due to increases in the probabilities of mid-banded (at sites with a relatively high number of dry days) and three-banded snails. The probability of an unbanded yellow snail decreases with increasing UHI. This is due to an increase in the probability of mid-banded and three-banded (but not five-banded). For pink snails, an increase in UHI increases the probability of a three-banded shell. The effect of temperature on banding types of pink snails seems to depend on the number of dry days at each location.

**Table 2.**
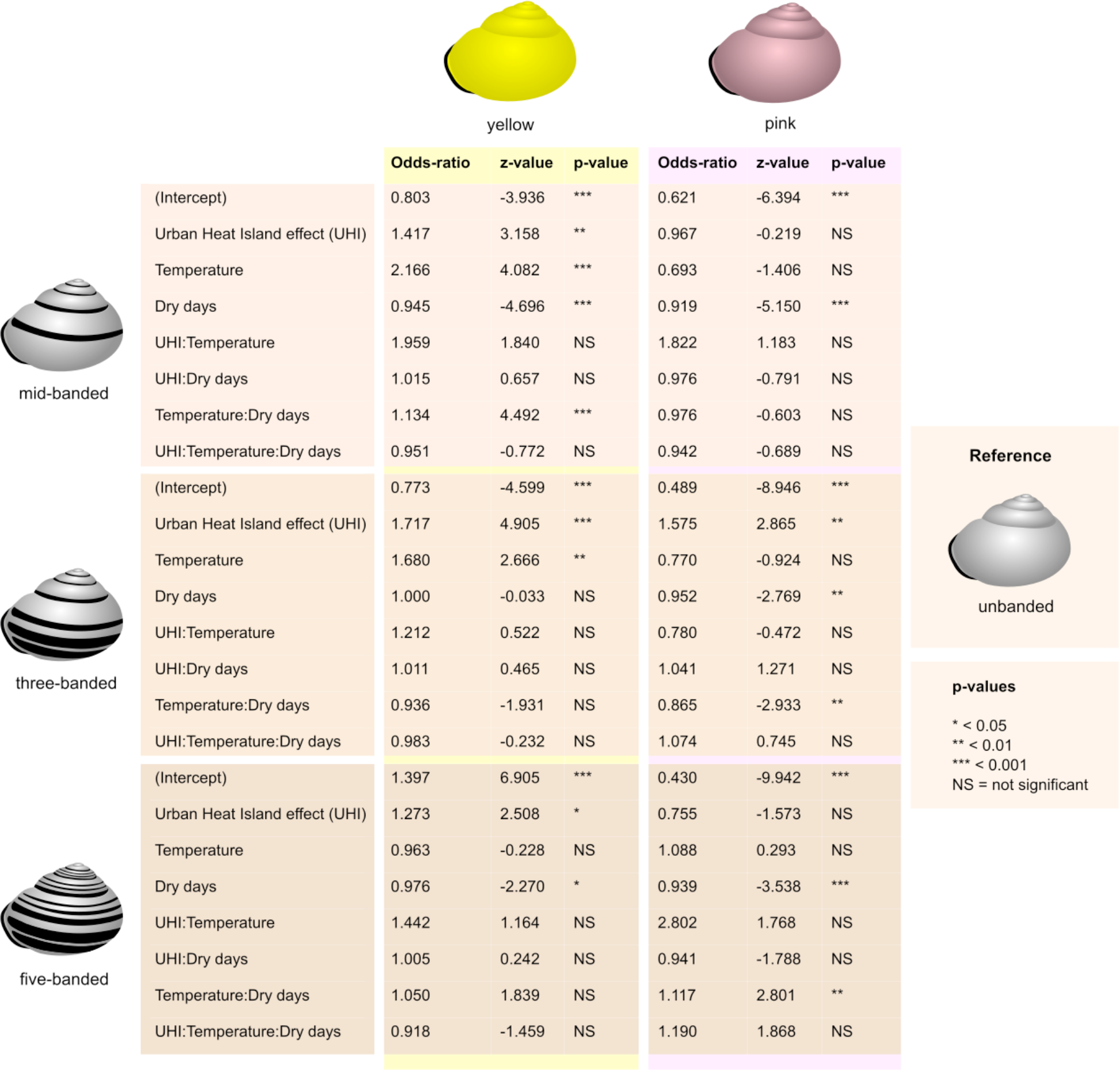
Results of multinomial regression models with banding type (unbanded, mid-banded, three-banded or five-banded) as outcome variable, for yellow (n = 3,660) and pink (n = 1,695) snails. Unbanded snails are the reference.

|  |  |  |  |  |  |  |  |  |  |
| --- | --- | --- | --- | --- | --- | --- | --- | --- | --- |
|  |  | yellow |  |  |  | pink |  |  |  |
|  |  | Odds-ratio | z-value | p-value | Odds-ratio | z-value | p-value |  |  |
| mid-banded | (Intercept) | 0.803 | -3.936 | *** | 0.621 | -6.394 | *** |  |  |
|  | Urban Heat Island effect (UHI) | 1.417 | 3.158 | ** | 0.967 | -0.219 | NS |  |  |
|  | Temperature | 2.166 | 4.082 | *** | 0.693 | -1.406 | NS |  |  |
|  | Dry days | 0.945 | -4.696 | *** | 0.919 | -5.150 | *** |  |  |
|  | UHI:Temperature | 1.959 | 1.840 | NS | 1.822 | 1.183 | NS |  |  |
|  | UHI:Dry days | 1.015 | 0.657 | NS | 0.976 | -0.791 | NS |  |  |
|  | Temperature:Dry days | 1.134 | 4.492 | *** | 0.976 | -0.603 | NS |  |  |
|  | UHI:Temperature:Dry days | 0.951 | -0.772 | NS | 0.942 | -0.689 | NS |  |  |
| three-banded | (Intercept) | 0.773 | -4.599 | *** | 0.489 | -8.946 | *** | unbanded | **Reference** |
|  | Urban Heat Island effect (UHI) | 1.717 | 4.905 | *** | 1.575 | 2.865 | ** |  |  |
|  | Temperature | 1.680 | 2.666 | ** | 0.770 | -0.924 | NS |  |  |
|  | Dry days | 1.000 | -0.033 | NS | 0.952 | -2.769 | ** |  |  |
|  | UHI:Temperature | 1.212 | 0.522 | NS | 0.780 | -0.472 | NS |  |  |
|  | UHI:Dry days | 1.011 | 0.465 | NS | 1.041 | 1.271 | NS |  |  |
|  | Temperature:Dry days | 0.936 | -1.931 | NS | 0.865 | -2.933 | ** |  |  |
|  | UHI:Temperature:Dry days | 0.983 | -0.232 | NS | 1.074 | 0.745 | NS |  |  |
| five-banded | (Intercept) | 1.397 | 6.905 | *** | 0.430 | -9.942 | *** | **p-values**<br>* < 0.05<br>** < 0.01<br>*** < 0.001<br>NS = not significant |  |
|  | Urban Heat Island effect (UHI) | 1.273 | 2.508 | * | 0.755 | -1.573 | NS |  |  |
|  | Temperature | 0.963 | -0.228 | NS | 1.088 | 0.293 | NS |  |  |
|  | Dry days | 0.976 | -2.270 | * | 0.939 | -3.538 | *** |  |  |
|  | UHI:Temperature | 1.442 | 1.164 | NS | 2.802 | 1.768 | NS |  |  |
|  | UHI:Dry days | 1.005 | 0.242 | NS | 0.941 | -1.788 | NS |  |  |
|  | Temperature:Dry days | 1.050 | 1.839 | NS | 1.117 | 2.801 | ** |  |  |
|  | UHI:Temperature:Dry days | 0.918 | -1.459 | NS | 1.190 | 1.868 | NS |  |  |

For both yellow, and in particular, pink snails, an increase in the number of dry days in general reduces the probability of a banded versus an unbanded shell. This indicates that, even though the number of dry days is not a significant predictor for the probability of a pink versus a yellow snail (table 1, figure 3a), the number of dry days does seem to influence pink snails: with increasing numbers of dry days, the proportion of unbanded snails becomes larger (table 2, figure 3b).

**Figure 3.**
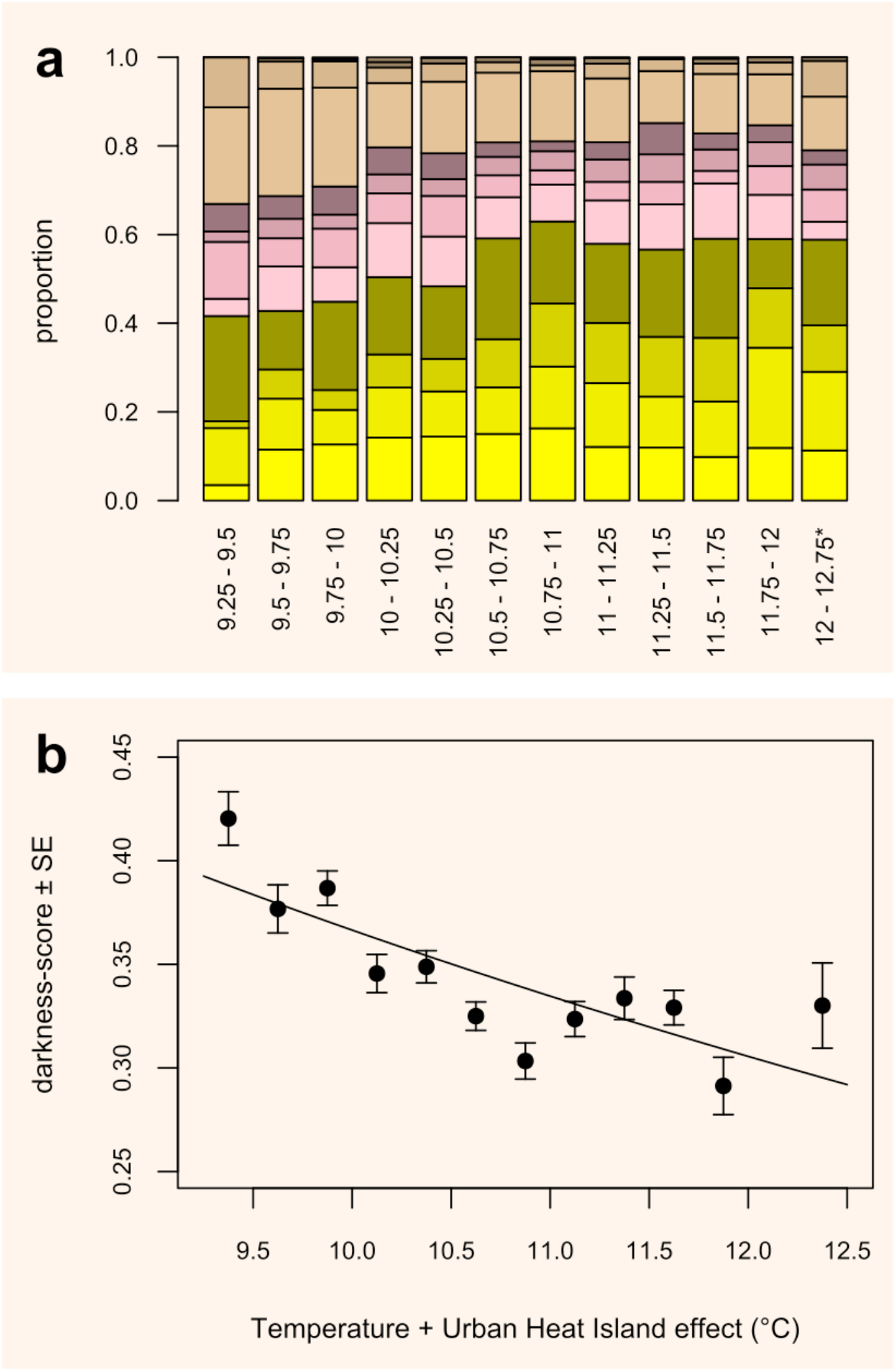
The proportion of colour morphs (a: yellow, pink and brown; per colour, from bottom to top: unbanded, midbanded, three-banded, five-banded; n = 6,809) and the average darkness-score (b; n = 6,838) per temperature+UHI category. Each category spans 0.25 °C, starting from 9.25°C, except the final category, which spans 0.75°C (three categories were grouped to attain a sample size comparable to the other categories). In b, the average darkness-score is plotted against the central value of each category. Error bars represent standard errors. There is a negative relationship between the value of temperature+UHI and the average darkness­score (darkness = 1.136 – 0.334*ln(Temperature+UHI); F = 18.09, p < 0.01, R^2^ = 0.64).

The combined effect of temperature and UHI is shown in figure 3. Temperature + UHI explains a high proportion (figure 5B, R^2^ = 0.64) of the variance in average darkness per temperature + UHI category; on average, darker snails were photographed at colder sites.

## Discussion

We used a citizen-science approach to obtain data on shell colour in Dutch urban and rural ***Cepaea nemoralis*** populations. As in the earlier, and comparable, “Evolution MegaLab"^27^, we used data on shell colour to reveal population-genetic patterns that may be related to human-induced environmental change. Our approach, however, differed from the “Evolution MegaLab” in various ways. First, rather than creating a stand-alone project, we adopted the social and digital network of an existing Dutch citizen science platform, Waarneming.nl (currently around 17,000 active users and 7,000,000 logged observations annually. Second, we minimized the effort required for adding data points: users could use a simple “point-and-shoot” smartphone app and did not need to supply any additional data. Finally, we specifically aimed at investigating urban evolution, by encouraging participants to record ***Cepaea nemoralis*** both inside and outside cities.

As shell colour and banding pattern in ***Cepaea*** are almost entirely genetically determined^24^, the differences in urban / non-urban shell coloration among the nearly 10,000 snails that were recorded can be interpreted in the context of urban evolution. We found that shells in urban environments are more likely to be yellow (and less likely to be pink) than in non-urban environments. The logistic regression shows that this is chiefly related to the urban heat island. These results suggest that urban conditions select for yellow snails and against pink snails. The effect is seen in both urban green and urban grey areas, which suggests that the main factor is overall urban air temperature, rather than structural or biological habitat features. There does seem to be a limit to this effect, since at the extreme temperature and UHI end, the proportion ceases to rise.

It is known from experimental work^8,14,18^ that yellow ***Cepaea nemoralis***, either because of a greater albedo or other causes, are better able to survive under high temperature regimes. It is therefore plausible that our results reveal natural selection imposed by the urban heat island on the (recessive) yellow allele for the shell ground colour gene in ***C. nemoralis***.

The results for shell banding are somewhat more complex. Dark spiral bands on the shell reduce a snail’s albedo^8^ and would therefore be expected to be selected against in the urban heat island. However, we found that this is only true for 5-banded morphs. The other main banding morphs (mid-banded and three-banded) in fact increase under urban conditions as well as under increasing temperature. Longitudinal studies of the evolutionary response of ***C. nemoralis*** to climate warming obtained similar results^16,27^. There are several possible explanations for this counter-intuitive effect. First of all, it is known that shell banding in ***C. nemoralis*** interacts with environmental factors such as shell-crushing predators: banded snails are stronger^20^ and, in certain habitats, better camouflaged^2,3^. Although we could not test for them, these evolutionary pressures may also differ in urban versus non-urban contexts. Furthermore, it is not impossible that the bands affect thermoregulation in yet underexplored ways. In the related ***Theba pisana*** it was found^13^ that dark bands on a light-coloured shell enhance cooling after an exposure to heat, possibly by creating differential airflow between dark and light areas, and by more rapid heat dissipation from the stripes. If such effects exist in ***C. nemoralis*** as well, they could combine with the greater albedo in yellow shells to select for yellow urban snails that have the exposed, top part of the shell unbanded (and therefore more reflective), while bearing bands on the underside (i.e., yellow mid- and three-banded) that may enhance cooling.

In summary, we show that a simple smartphone app, linked to a citizen science web platform, enables effective monitoring of phenotypic change in the urban heat island, probably as a result of natural selection. Our results concern the data collected in the first year, but the app continues to generate data, which could be used to confirm our results in future years.

The system we describe could also be expanded to other organisms that are known to display easily observable phenotypic evolution in urban contexts, such as plumage coloration in rock pigeons, ***Columba livia***^5,6^, and male “necktie” width in the great tit, ***Parus major***^25^. This would allow the development of a continuous monitoring system of urban evolution in multiple organisms — especially if current attempts to replace human validation with artificial intelligence in citizen science apps^21^ are successful.

## Methods

### SnailSnap

A smartphone application ("SnailSnap") was developed for Android-devices. The app is available for download, free of charge, at Google Play Store (https://play.google.com/store/apps/details?id=nl.zostera.slakkenschieter). It was designed as a convenient and accessible method to upload images of live, adult ***Cepaea nemoralis*** to Waarneming.nl (the Dutch version of Observation.org). The SnailSnap project officially began on April 1^st^, 2017. Prior to this date, we sent out a press release. We also developed a website (www.snailsnap.nl), gave interviews on national radio, and handed out flyers at nature-related events. During the entire course of the project, we used the public social media channels (Facebook and Twitter) of the Netherlands natural history museum Naturalis, and the online newsletters of Waarneming.nl and insect knowledge centre EIS to inform the general public.

Participants were instructed (via the website and instructions incorporated into the app) on how to find, recognize, and photograph adult ***C. nemoralis***. They were asked to take a single photograph per snail, to include only one snail per picture, and to take the pictures at such an angle that the lip and all bands on the shell were visible. Images and the associated GPS-coordinates were automatically uploaded to the Waarneming.nl server as soon as the participant’s smartphone connected to a wifi-network. Alternatively, app-users could send the pictures to the server via mobile internet using the Send-button that was incorporated in the app.

Participants could also upload pictures to the server via the Waarneming.nl website and its apps iObs and Obsmapp. All pictures that were uploaded were then handled by a group of ten validators, who (i) checked the correct identification as ***C. nemoralis***, and (ii) attributed the correct colour and banding code to each snail. All snails with a visible dark lip were included in the data set. ***Cepaea hortensis***, which can be distinguished from ***C. nemoralis*** by its white lip, is rare in most parts of the Netherlands. Therefore, only those pictures without a visible dark lip were removed from the data set if they were taken in one of three known areas where ***C. hortensis*** is relatively common (namely, the south of the province of Limburg [latitude < 51.2], the northeast of the province of Groningen [latitude > 53, longitude > 6.5] and the area east and north-east of Nijmegen [51.8 < latitude < 52, longitude > 5.75]).

We chose October 15^th^, 2017, as end date (in the Netherlands, ***C. nemoralis*** starts hibernating around this date). Not all uploaded images could be classified to colour morph. Some were very unclear or had been taken from an inappropriate angle. Sometimes single snails were photographed more than once (in this case, only one of the pictures was classified). Quite a large number of pictures had been taken from an apical angle, hence the lower bands (4^th^ and 5^th^) were not visible. Snails on such pictures were classified into a special group of codes and used only for analyses of shell ground colour (not banding). The remaining observations were relatively evenly distributed over the country (figure S2).

### Accuracy of the SnailSnap data

To test the accuracy of the colour and banding classification by validators, we double-checked 240 snails (60 snails starting at four random dates, i.e., 1^st^ of August, 15^th^ of July, 18^th^ of September, and 28^th^ of April). We found that the banding pattern was only misclassified once. In 33 out of the 240 cases, however, our colour classification did not match the classification by the validator. The most common discrepancies were dark pink snails classified as brown, and multi-banded pink snails as yellow. All validators had been given instructions on how to classify snails. Also, during the course of the project, validators who occasionally misclassified snail colours had been asked to revisit the instruction documents and/or to adjust the settings of their computer screen. Furthermore, in several cases probably we were the ones to misclassify a snail. Mainly due to the variation within each of the three main colour categories, assigning a colour to a snail is not always straightforward. This is especially true for people who are not experienced with studying ***Cepaea***. Although it is clear that the current colour classification method is not perfect, it is probably more accurate than the method used in the “Evolution Megalab", a previous citizen science project on ***Cepaea***^27^.

During the course of the project, we noticed that consecutive pictures taken in the app by the same user sometimes had the same GPS-coordinates. It transpired that certain conditions (a low quality mobile phone, cloudy weather, tall buildings) could cause a GPS-fix to fail. Pictures would then be associated with the last successful GPS-fix until a new GPS-fix was made. Although this problem is, to some extent, unavoidable (highly accurate GPS-fixes could take unacceptably long under some conditions), the app was updated to improve this condition on September 11. We have no way of knowing to what extent this GPS-problem has influenced the accuracy of locality data for the observations, but we expect it to be negligible.

### Data analysis

All data on shell colour, banding, and location were exported from the Waarneming.nl server to a spreadsheet. The 7,868 individual data points, each representing a single snail, were then imported into an ArcGIS v10.2.2 (Environmental Systems Research Institute, Redlands, CA) project. Using the tool “Extract Multi Values to Points” and the “Join data” dialog box (join by location), environmental and climatic geographical data were added to the data points based on the location of each point. We used geographical data on the Urban Heat Island effect (°C, grid, 10×10 meters), the average yearly temperature (°C, grid, 1000 × 1000 m), the average number of dry days per year (grid, 1000 × 1000 m) and habitat types (land use, vector) (see table S2 for more details).

All statistical analyses were performed in R (version 3.4.3 [Ref. 17]). To compare proportions of colour morphs from different habitats, we included only snails with a complete banding code that fit into one of the four main banding types (unbanded, midbanded, three-banded, and five-banded; 6,809 in total). Based on the GPS-coordinates, snails were assigned to one of four habitat types: (1) agricultural land (n = 832), (2) nature (including forests; n = 888), (3) urban green areas (urban parks and forests, sport and recreational areas; n = 817) and (4) urban “grey” areas (residential, commercial, and industrial areas; n = 4,008). The proportions of snails of different colours and/or banding types were compared between habitat types using Chi-squared tests.

Multinomial logistic regression models with shell colour and banding type as outcome variable were run using the “multinom” function from the nett package^28^. Odds-ratios, z-values and p-values were obtained via the “tidy” function from the broom package^19^. All predictor variables were centred around zero (centred X = X – mean(X); centred predictor values were calculated for each model separately). To interpret marginal effects and significant interaction terms, effects were displayed using the effects package^7^.

To visualize the combined effect of temperature and UHI (temperature + UHI), this variable was divided into categories and proportions of colour morphs were calculated per category.

To study the combined effect of shell colour and banding patterns, for each snail with a complete banding code (n = 6,838), a score was calculated based on the known thermal properties of colour morphs^8^. Using yellow unbanded snails as the baseline, the expected internal temperature increases of the other morphs were calculated by adding 0.3°C for pink, 0.6°C for brown, 0.07°C for each band and 0.03°C for each band fusion. The resultant expected temperature increase was used as “darkness score"^22^. For temperature + UHI, average darkness scores were calculated per category. These average darkness scores were plotted against the central variable value of each category and a regression line was fitted.

To illustrate the interaction between temperature and UHI, colour morph proportions were calculated per UHI-category for snails twice: once for sites with an average temperature below or equal to 10°C (the median temperature) and once for sites with an average temperate above 10°C.

## Acknowledgements

We gratefully acknowledge all users of the app and other participants, especially school classes and their teachers. We especially thank the validators, Marleen Baltus, Mascha Castelijns-Dolne, Gijs van den Ende, Michael Inden, Jesse Kerkvliet, and Koen Verhoogt, for investing time in checking identifications. Staff and volunteers at Insect Knowledge Centre EIS supported this project throughout. Apps were developed by Zostera. Waarneming.nl hosted our project and facilitated data transfer. The citizen science component was developed in close collaboration with the department for Education at Naturalis Biodiversity Center. Jésus Aguirre and Tom van Dooren provided helpful advice on the statistical analysis of GIS data. Financial support came from the Gratama Foundation and the Van der Hucht De Beukelaar Foundation.

## Author contributions

M.S. and V.K. conceived the study. T.B., M.S. & V.K. designed the set-up of the app and the citizen-scientists’ involvement. T.B. supervised the testing and improvement of the app, N.A.G.K. supervised the data collection and their validation. N.A.G.K. analysed the data. N.A.G.K. & M.S. wrote the paper.

## References

1. Alberti, M. et al. Proc. Natl. Acad. Sci. USA 114, 8951–8956 (2017).

2. Cain, A. J., & Sheppard, P. M. Heredity 4, 275–294 (1950).

3. Cain, A. J., & Sheppard, P. M. Genetics, 39, 89–116 (1954).

4. Chandler, T. J. The Climates of London (Hutchinson, London, 1965).

5. Chatelain, M., Gasparini, J., Jacquin, L., & Frantz, A. Biol. Lett. 10, 20140164 (2014).

6. Chatelain, M., Gasparini, J., & Frantz, A. Global Change Biol. 22, 2380–2391 (2016).

7. Fox, J. & Hong, J. J. Stat. Software 32, 1–24 (2009).

8. Heath, D. J. Oecologia, 19, 29–38 (1975).

9. Hendry, A. P., Gotanda, K. M., & Svensson, E. I. (2017). Phil. Trans. R. Soc. B, 372, 20160028 (2016).

10. Johnson, M. T., Thompson, K. A., & Saini, H. S. Am. J. Bot. 102, 1951–1953 (2015).

11. Johnson, M. T., & Munshi-South, J. Science 358, eaam8327 (2017).

12. Jones, J. S., Leith, B. H., & Rawlings, P. Annu. Rev. Ecol. Syst. 8, 109–143 (1977).

13. Knigge, T., Di Lellis, M. A., Monsinjon, T., & Köhler, H. R. J. Therm. Biol. 69, 54–63 (2017).

14. Lamotte, M. Cold Spring Harbor Symp. Quant. Biol. 24, 65–86 (1959).

15. Ożgo, M. J. Moll. Stud. 80, 286–290 (2014).

16. Ożgo, M., & Schilthuizen, M. Global Change Biol. 18, 74–81 (2012).

17. R Core Team R: A language and environment for statistical computing (R Foundation for Statistical Computing, Vienna, 2017).

18. Richardson, A. M. Nature 247, 572 (1974).

19. Robinson, D. broom: Convert Statistical Analysis Objects into Tidy Data Frames. R package version 0.4.4 (https://CRAN.R-project.org/package=broom, 2018).

20. Rosin, Z. M., Kobak, J., Lesicki, A., & Tryjanowski, P. Naturwissenschaften 100, 843–851 (2013).

21. Schermer, M., & Hogeweg, L. Biodiv. Inf. Sci. Stand. 2, e25268 (2018).

22. Schilthuizen, M. Heredity 110, 247–252 (2013).

23. Schilthuizen, M. Darwin Comes to Town: How the Urban Jungle Drives Evolution (Quercus, London, 2018).

24. Schilthuizen, M., & Kellermann, V. Evol. Appl. 7, 56–67 (2014).

25. Senar, J. C., Conroy, M. J., Quesada, J., & Mateos-Gonzalez, F. Ecol. Evol. 4, 2625–2632 (2014).

26. Silvertown, J. Trends Ecol. Evol. 24, 467–471 (2009).

27. Silvertown, J. et al. PLoS One 6, e18927 (2011).

28. Venables, W. N. & Ripley, B. D. Modern Applied Statistics with S. Fourth Edition (Springer, New York, 2002).

29. Worthington, J. P. et al. Meth. Ecol. Evol. 3, 303–309 (2012).

